# Dissecting the *Pyrenophora tritici-repentis* (tan spot of wheat) pangenome

**DOI:** 10.1101/2022.03.07.483352

**Authors:** Ryan Gourlie, Megan McDonald, Mohamed Hafez, Rodrigo Ortega-Polo, Kristin E. Low, D. Wade Abbott, Stephen E. Strelkov, Fouad Daayf, Reem Aboukhaddour

## Abstract

We sequenced the genome of a global collection (40 isolates) of the fungus *Pyrenophora tritici-repentis* (Ptr), a major foliar pathogen of wheat and model for the evolution of necrotrophic pathogens. Ptr exhibited an open-pangenome, with 43% of genes in the core set and 57% defined as accessory (present in only a subset of isolates), of which 56% were singleton genes (present in only one isolate). A clear distinction between pathogenic and non-pathogenic genomes was observed in size, gene content, and phylogenetic relatedness. Chromosomal rearrangements and structural organization, specifically around the effector coding genes, were explored further using the annotated genomes of two isolates sequenced by PacBio RS II and Illumina HiSeq. The Ptr genome exhibited major chromosomal rearrangements, including chromosomal fusion, translocation, and segment duplications. An intraspecies translocation of *ToxA*, the necrosis-inducing effector-coding gene, was facilitated within Ptr via a 143 kb ‘*Starship’* transposon (dubbed ‘Horizon’). Additionally, *ToxB*, the gene encoding the chlorosis-inducing effector, was clustered as three copies on a 294 kb transposable element in a ToxB-producing isolate. *ToxB* and its carrying transposon were missing from the *ToxB* non-coding reference isolate, but the homolog *toxb* and the transposon were both present in another non-coding isolate. The Ptr genome also appears to exhibit a ‘one-compartment’ organization, but may still possess a ‘two-speed genome’ that is facilitated by copy-number variation as reported in other fungal pathosystems.

**IMPORTANCE:** Ptr is one of the most destructive wheat pathogens worldwide. Its genome is a mosaic of present and absent effectors, and serves as a model for examining the evolutionary processes behind the acquisition of virulence in necrotrophs and disease emergence. In this work, we took advantage of a diverse collection of pathogenic Ptr isolates with different global origins and applied short- and long-read sequencing technologies to dissect the Ptr genome. This study provides comprehensive insights into the Ptr genome and highlights its structural organization as an open pangenome with ‘one-compartment’. In addition, we identified the potential involvement of transposable elements in genome expansion and the movement of virulence factors. The ability of effector-coding genes to shuffle across chromosomes on large transposons was illustrated by the intraspecies translocation of *ToxA* and the multi-copy *ToxB*. In terms of gene contents, the Ptr genome exhibits a large percentage of orphan genes, particularly in non-pathogenic or weakly-virulent isolates.

## INTRODUCTION

*Pyrenophora tritici-repentis* (Ptr) is an ascomycete fungus that causes tan spot, one of the most destructive foliar diseases of wheat worldwide (1, 2). The Ptr genome represents an exciting case to explore the evolution of virulence and the emergence of new diseases. Ptr has served as a model species for necrotrophic plant pathogens, as it has the ability to produce various necrotrophic effectors, previously known as host-selective toxins (2). To date, three effectors have been identified and are known virulence factors: ToxA is a necrosis-inducing effector, while ToxB and ToxC are chlorosis-inducing effectors (3). Additional effectors may be involved in pathogenicity and await characterization (4). ToxA and ToxB are ribosomally-synthesized proteins, and while ToxA is encoded by a single copy *ToxA* gene, ToxB is encoded by the multi-copy *ToxB* genes (3). Understanding of ToxC-coding gene(s) and its synthesis awaits further characterization (5, 6). In Ptr, the *ToxA* gene is present only in isolates that secrete the ToxA effector, and no homolog has been identified in non-producing isolates of Ptr. In contrast, *b* homologs, termed *toxb*, are present in some Ptr isolates that do not secrete the active form of the ToxB protein. Homologs of both *ToxA* and *ToxB* have been reported in related and distantly related fungal species (7–10).

Ptr isolates are grouped into eight different races (races 1 to 8) based on which combinations of effectors they secrete (3). In North America and Australia, Ptr isolates are mainly ToxA and ToxC-producers, with ToxB-producers being extremely rare or absent (4, 11). In regions encompassing the wheat centre of origin, like the Fertile Crescent and North Africa, all races and effectors are found even in under-surveyed regions (2, 4). ToxB-producing isolates were found to predominate in North Africa (4, 11, 12). The lack of isolates with *ToxB* in North America and Australia indicates divergent evolution of Ptr and its effectors worldwide. An independent origin of these effectors was suggested previously (7, 8, 13, 14). For example, *ToxA* was acquired by horizontal-gene-transfer (HGT) from a related fungal species (7, 13, 14), while *ToxB* (and its homologs) are believed to have evolved via vertical inheritance (7, 8, 13, 15).

The Ptr genome has been described as highly plastic, with evidence of major chromosomal rearrangements, including insertions, deletions, inversions, and translocations (13, 14, 16–18). The first full Ptr genome was released by the Broad Institute in 2007, based on a race 1 isolate (Pt-1C-BFP, abbreviated throughout as BFP) from the United States, and was assembled to the chromosome level with the aid of optical mapping (16). In addition, the full genomes of 11 Ptr isolates from North America and Australia have also been sequenced and are now publicly available (16, 17, 19). Of those, three (M4, V1, and DW5) were assembled from long-read sequences.

Previously, DNA transposons and long terminal repeat (LTR) retrotransposons were associated with genome-wide segmental duplications, insertions, and chromosomal fusions in Ptr (15). However, genomic regions with AT-rich content, which are associated with repeat-induced point mutations (RIP), were not reported at high frequencies in previously sequenced Ptr isolates from Australia and the USA (17). RIP is often associated with the “two-speed genome” model in a number of fungal pathogens; this model describes the pattern of physical genome compartmentation with repeat-rich gene-sparse regions and repeat-sparse gene-dense regions (20). Here, we sequenced 40 Ptr isolates of diverse origins and races, and performed a whole pangenome analysis with the addition of the reference isolate (BFP), for a total of 41 isolates. These isolates represent all known races including a newly identified North African atypical race that induces necrosis but lacks the *ToxA* gene (4). In addition to the pangenome, we provide a detailed look at gene contents, phylogenetic relatedness, and effector gene movement. In this study, polished long-read assemblies representing the full genome of two isolates (I-73-1 and D308) were explored to define the structural organization and compartmentation of the Ptr genome by comparison with three previously published genomes (BFP, DW-5, and M4). I-73-1 is a race 8 isolate collected from Syria and represents the most complex race, producing all three known effectors (ToxA, ToxB, and ToxC). D308 is a race 3 isolate collected from Canada, and codes for ToxC only. This work provides a detailed comparative genomic analysis of Ptr for use by the research community on fungal pathogens.

## RESULTS

### Hybrid assemblies capture a better estimate for genome size

The *de novo* assemblies of Illumina short-reads indicated a variable genome size ranging from 34.1 Mb to 36.9 Mb, with an average of 34.82 Mb. The smallest assembly was for isolate T128-1 (34.12 Mb), which has been identified as an atypical isolate with a novel virulence phenotype (4). The largest assemblies were for isolates 92-171R5 (36.8 Mb, race 5) and G9-4 (36.97 Mb, race 4).

The hybrid assembly (PacBio RS II and Illumina Hi-Seq data) of isolate I-73-1 contained 39 contiguous sequences, with 11 chromosome-sized contigs greater than 2 Mb. Similar results were obtained with isolate D308, whose assembly contained 70 contiguous sequences, and 11 contigs greater than 1 Mb with an additional two smaller contigs representing the majority of the genome. Genome completeness was assessed by Benchmarking Universal Single-Copy Orthologs (BUSCO) using a set of conserved fungal genes, and all isolates (except G9-4) were assessed as >99% complete. The short-read assemblies of I-73-1 and D308 were ∼5.3 Mb smaller than the long-read-based assemblies. Summary statistics for all assemblies are available in Table 1.

**Table 1.**
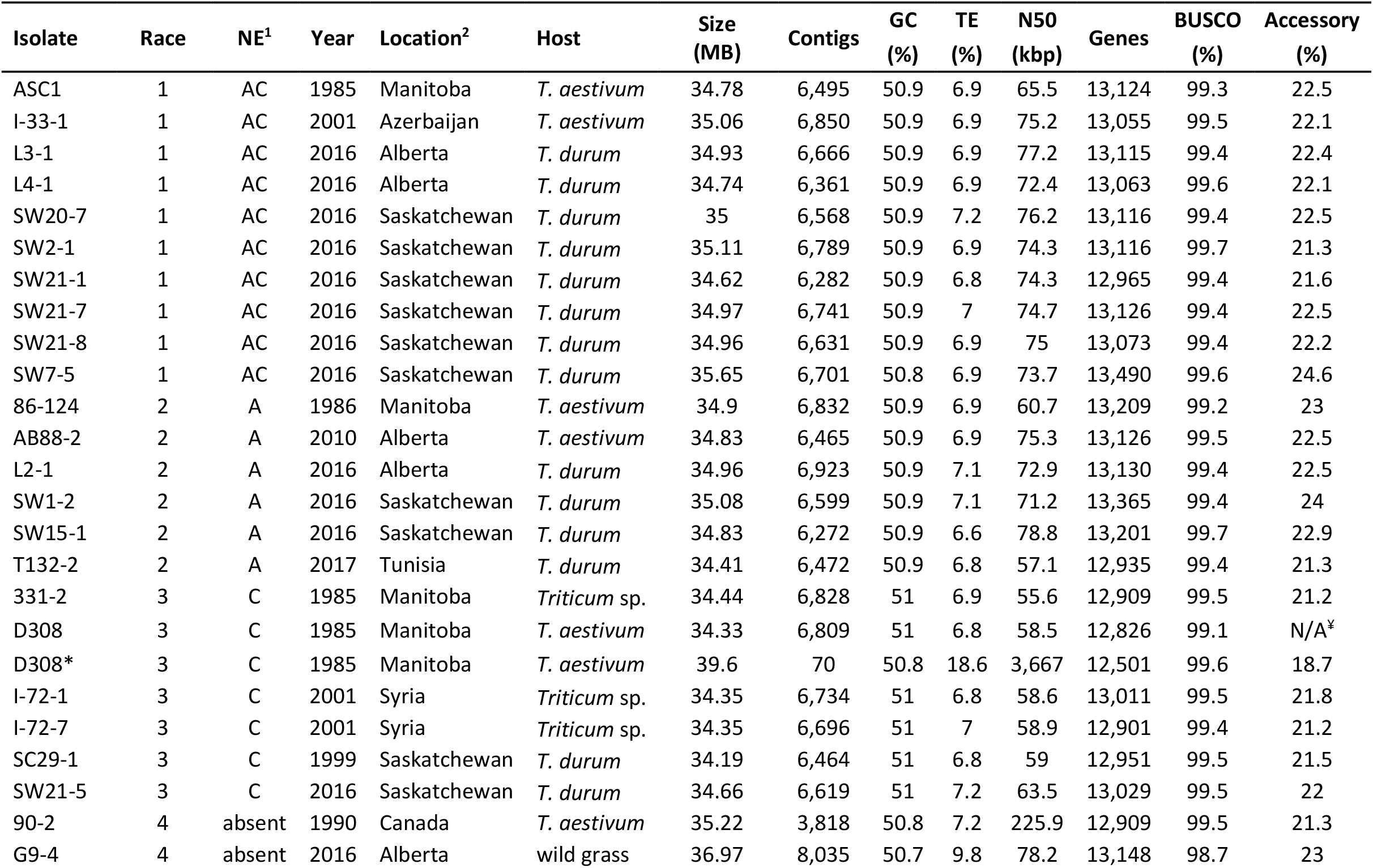
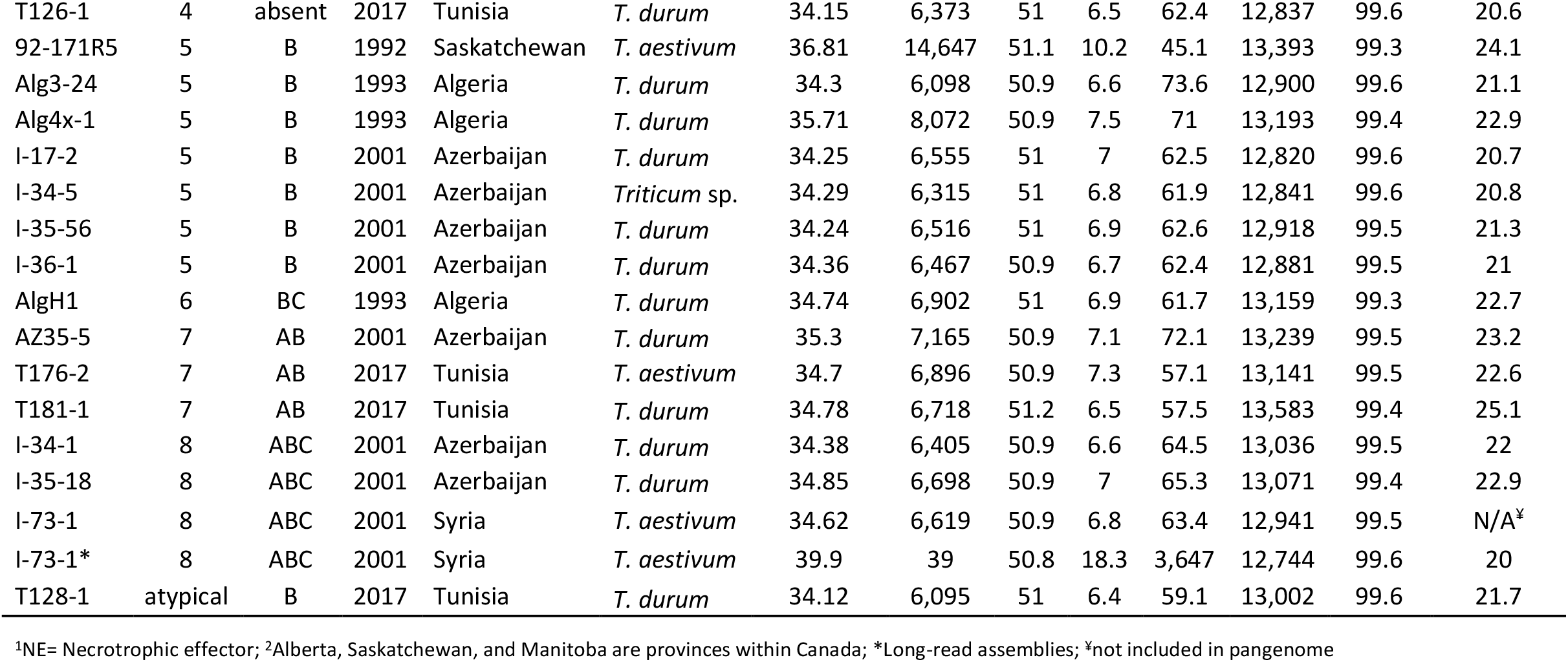
Summary statistics for 40 isolates of *Pyrenophora tritici-repentis* sequenced with Illumina HiSeq X and assembled with Shovill/SPAdes, and two isolates sequenced with PacBio and assembled with Flye/Pilon (table is sorted by race then isolate name).

### Ptr has an open pangenome with high accessory gene content and unique genes for non-pathogenic and weakly virulent isolates

On average, each isolate contained 13,071 predicted genes, and collectively, 522,848 predicted genes across all isolates were reported. After grouping by similarity, with a threshold of ≥90% (21), we were left with 23,454 unique gene clusters. In total, 10,159 genes were identified as core (conserved across all isolates) (43% of total pangenome) (Figure 1A). The remaining 13,295 gene models (∼57% of the total pangenome) were the accessory genome, with individual accessory sizes ranging from 2,661 to 3,424 genes. Within the accessory genome were 7,476 singletons (genes present in only one isolate) (56% of accessory genome; 32% of the total pangenome). Within the singletons, a large number 3,293 (44%) were present in isolates in the outgroup of the phylogeny (Figure 2A), including 90-2 (461 genes), 92-171R5 (1,598 genes), and G9-4 (1,234 genes). Additionally, isolate T128-1, which exhibits a novel necrotic phenotype (4), contained 54 unique genes.

**Figure 1.**
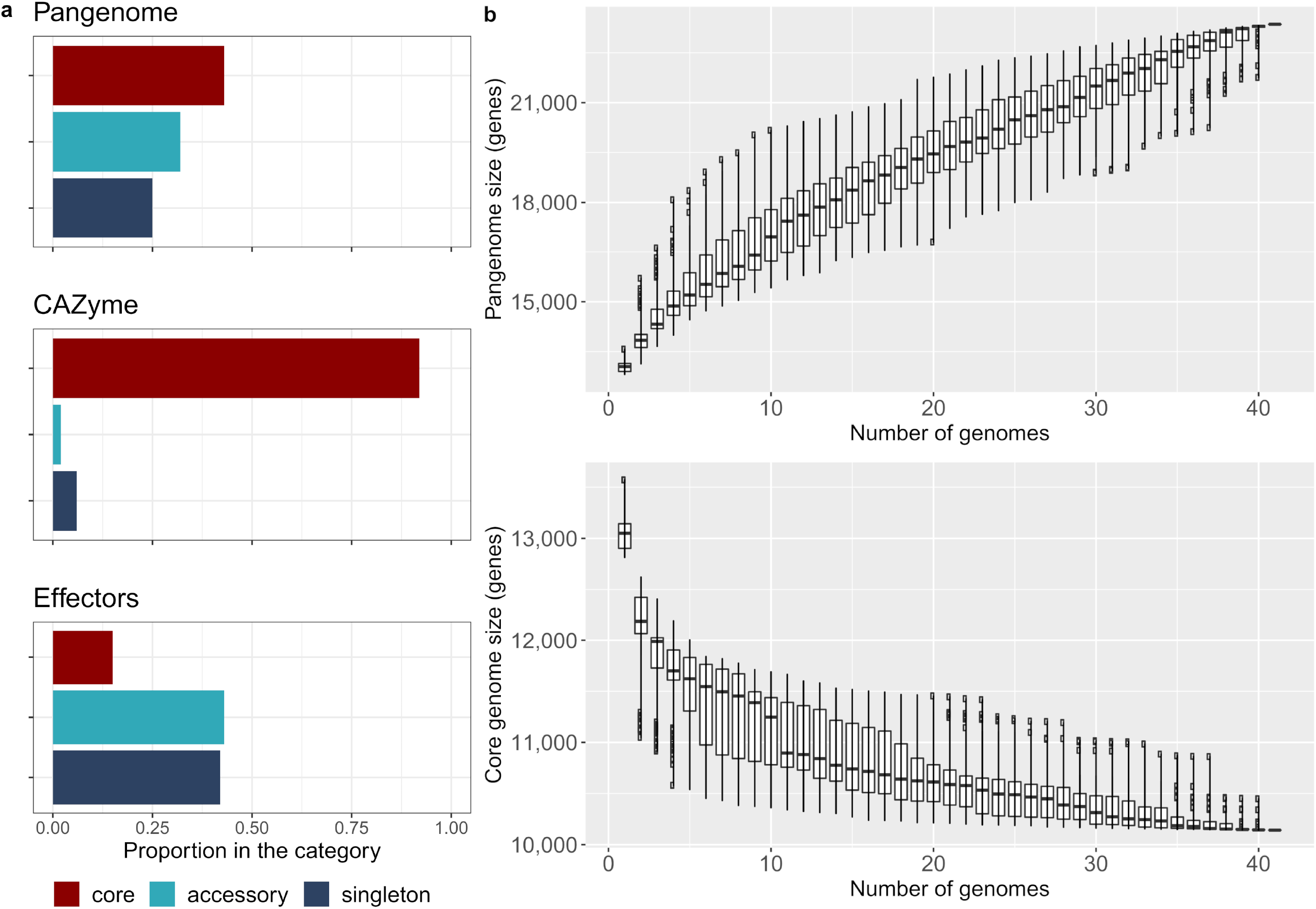
The pangenome of *Pyrenophora tritici-repentis*. **a** Proportion of genes present in core, accessory (excluding singletons), and singleton sets for the pangenome, CAZymes, and effectors. **b** Total number of unique genes and core genes in the pangenome of *Pyrenophora tritici-repentis* as genomes are added. Starting genome was always Pt-1C-BFP with subsequent genomes added

**Figure 2.**
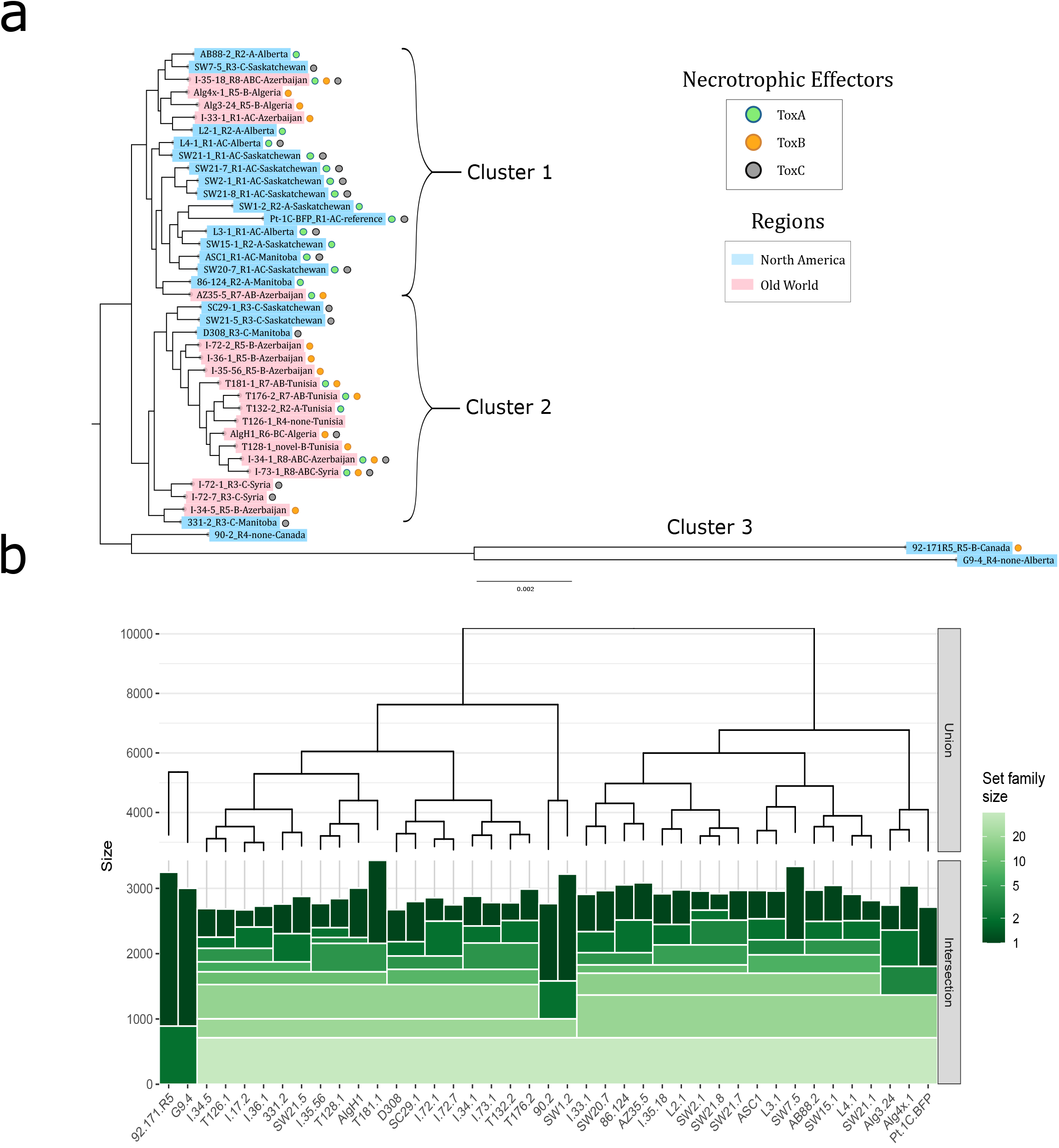
Evolutionary relatioships of isolates in the Ptr pangenome. **a** Maximum-likelihood phylogenetic tree of *Pyrenophora tritici-repentis* base on aligned and concatenated proteins present in the core genome (10,159 genes). Isolate names are followed by their race (R1 to R8) designation, necrotrophic effector(s) produced (A, B, and/or C), and their location of origin (isolates collected within Canada show province names; isolates outside Canada show country of origin). Presence of ToxA (blue circles) and ToxB (orange circles) confirmed via BLAST, and presence of ToxC (grey circles) determined by phenotyping. Isolates were color coded by region with North America representing Canada and the United States and Old World representing Caucuses, North Africa, and Fertile Crescent. **b** Hierarchical sets of *Pyrenophora tritici-repentis* accessory pangenome, size represents number of genes shared by isolates in each cluster, where inclusion in a cluster is based on any given isolate overlapping with the horizontal bars.

Genes were binned into ‘clusters’ based on the presence or absence of genes to provide insights into gene gains and losses in Ptr (Supplemental Figure 1). Each cluster is indicative of how many isolates share a particular gene (e.g., cluster 2 shows genes shared by two isolates, cluster 3 by three isolates, etc.). Clusters 1 and 41 were not included, as these represent a singleton and the core gene set, respectively. The result of this clustering is groups of genes (grouped by how many isolates contain the gene) many of which are likely gene gains and losses throughout Ptr.

Isolates carrying the *ToxA* gene had significantly larger accessory genomes (mean of 2,987 genes) compared with isolates lacking *ToxA* (mean of 2,820 genes) based on a two-sample t-test (*p* = 0.002). No such relationship was found with *ToxB* or putative *ToxC-*carrying isolates. An ANOVA followed by Tukey’s HSD test indicated that race 7 (*ToxA, ToxB*) and race 3 (*ToxC*) isolates also had significantly different accessory genome sizes (average of 3,162 and 2,779 genes, respectively; *p* = 0.04), with no other notable differences between races. Additionally, the total number of genes present in the pangenome increased steadily with each additional genome (Figure 1B), and the number of core genes appears to have reached a stable level after the addition of 38 genomes (Figure 1B). The continual increase of genes per genome and the large accessory genome (57%) are strong indications that Ptr has an open pangenome.

### Ptr core protein and SNP phylogenies exhibit clustering based on necrotrophic effector secretion

A maximum-likelihood tree to estimate genetic relatedness among Ptr isolates was constructed using an alignment of the 10,159 core proteins, and showed that most isolates partitioned into two major clusters (Figure 2A). Cluster 1 contained 20 isolates, most of which were ToxA-producing Canadian isolates classified as races 1 and 2. Few isolates in cluster 1 were ToxB-producers; those that were ToxB-producers were from Azerbaijan and Algeria. Cluster 2 contained 19 isolates, most of which were ToxA non-producing races 3 and 5, which mostly originated in the Fertile Crescent and nearby regions. The Canadian isolates in cluster 2 were all classified as race 3 (ToxC producers). Three isolates (92-171R5, 90-2, and G9-4) clustered as an outgroup. 92-171R5 is the only race 5 isolate identified in Canada, and was described as weakly virulent (22). Isolates 90-2 and G9-4 were classified as non-pathogenic race 4 (12, 23).

Heirarchical set groupings based on accessory proteins revealed similar clustering as the core protein phylogeny (Figure 2B), with the only exceptions being in which major cluster the isolates 90-2 (race 4) and SW1-2 (race 1) were placed. These two isolates shared a high number of genes not present in other isolates, defining their placement within the hierarchical set. Clusters in the hierarchical sets were defined by the presence or absence of accessory genes and does not take into account any sequence variation in the homologs used to construct the protein phylogeny, but highlights how gene gains and losses can mirror mutation rates. Additional phylogenetic trees based on SNP data also closely match the core protein phylogeny and the hierarchical set trees in terms of isolate clustering, with the divergent isolates separating even further (Supplemental Figure 2A and 2B’).

### Predicted genes and effectors

Of the 23,454 unique proteins in the Ptr pangenome, only 46% were present in the Pfam functional database. Of the 10,159 core genes, 6,951 (69%) had some described functionality (e.g., domain, family, etc.). However, of the 13,295 accessory genes, only 3,782 (29%) were categorized, the remaining being hypothetical proteins. Core gene protein lengths were significantly greater than accessory proteins, with mean lengths of 482 aa and 241 aa, respectively, and medians of 411 aa and 156 aa. Singleton proteins were marginally smaller than other accessory proteins, with a mean length of 232 aa (median 156 aa). A small number of proteins (108) were quite large, with lengths exceeding 2,000 aa, and six of those exceeding 5,000 aa; the largest protein was 9,750 aa in length. Many of these genes were annotated in previously published Ptr genomes.

Effectors play a major role in establishing infection in necrotrophic pathogens. In total, 729 potential effectors (3.1% of the total pangenome) were predicted in Ptr. Of these, 106 (14.5%) were present as core genes, and 626 (85%) were accessory genes, with 313 of the accessories being singletons (Figure 1A). Omission of the non-pathogenic race 4 isolates (90-2, G9-4, and T126-1) revealed 584 unique effectors, with 127 genes in the core and 457 as accessory (198 singletons). Nearly half of the potential effectors were singletons present in the weakly virulent isolate 92-171R5.

### CAZymes

We identified 305 unique CAZymes from 91 CAZyme families that are known to be active on cell wall polysaccharides. Of these, 281 (92%) were observed in all isolates; the number of proteins within the CAZyme families were predominantly conserved across all isolates (Supplemental File 4). Although numbers were similar between isolates and were related at the sequence level, there may be functional differences that remain to be explored. CAZyme families associated with plant cell wall degradation and fungal phytopathogenesis were abundant (Supplemental File 1). These included GH5 cellulases and endoglucanases, GH43 arabinases and xylosidases, AA9 lytic cellulose monooxygenases, CE1 hemicellulases, and CE5 cutinases (24–29). Protein sequences within these families accounted for 31% of CAZymes identified. It should be noted that not all CAZymes are associated with pathogenesis, and many are likely involved in routine cellular activities, such as fungal cell wall remodeling and glycoprotein maturation.

### Chromosomal structural organization

Full genome alignment between the hybrid assemblies of isolate I-73-1 (race 8) and the reference BFP (race 1) showed 11 contigs in I-73-1 of comparable size and content to the 11 chromosomes of the reference BFP. Alignment of D308 (race 3) indicated 10 contigs of comparable size, with an additional three smaller contigs completing a full alignment to BFP. BFP chromosomes 4 and 7 appeared to be homologous between all three isolates (contigs 12 and 9 in I-73-1 and contigs 5, 11, and 14 in D308) (Figure 3). Chromosomes 3, 5, 6, and 10 in BFP were also mostly homologous with contigs 2, 7, 17 and 6, respectively, in I-73-1 (Figure 3); the only notable exception was the *ToxA* containing region detailed further below (Figure 4A).

**Figure 3.**
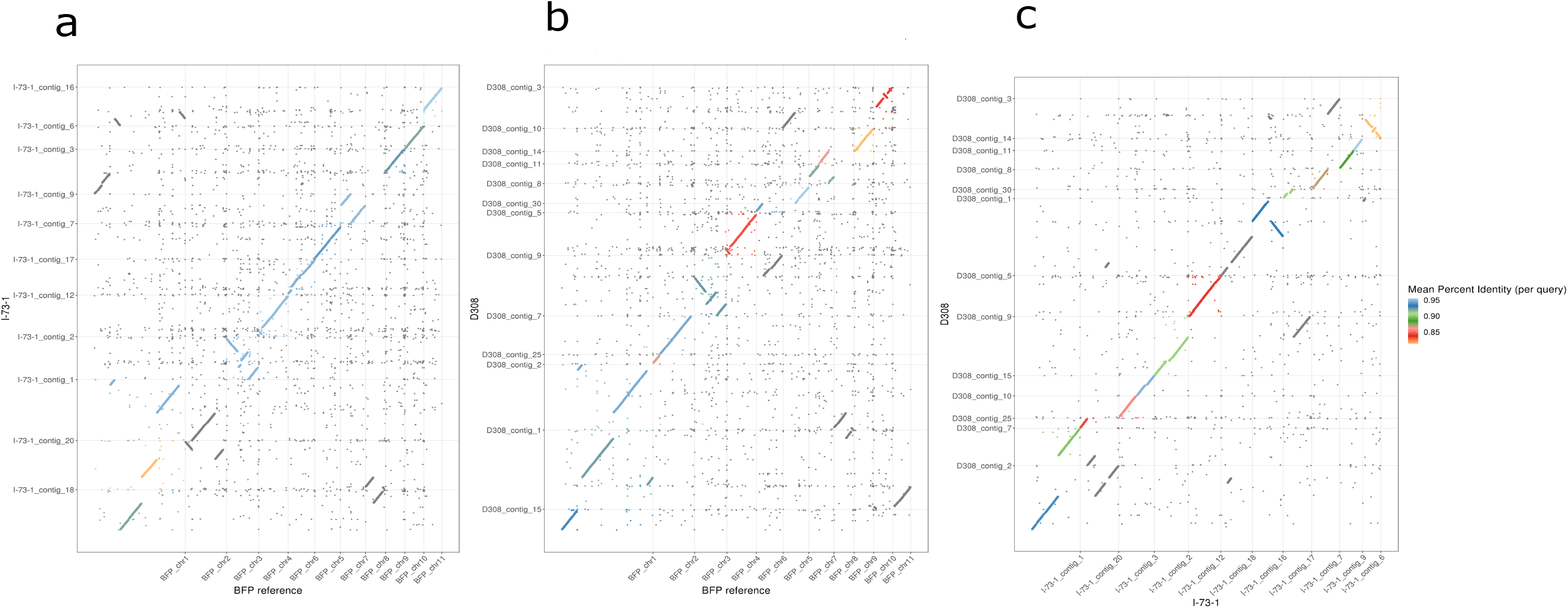
Full genome dotplot alignments of newly sequenced long-read assemblies of *Pyrenophora tritici-repentis* isolates I-73-1 (race 8) and D308 (race 3) to the reference isolate Pt-1C-BFP (race 1). **a** Pt-1C-BFP and I-73-1; **b** Pt-1C-BFP and D308; **c** I-73-1 and D308.

**Figure 4.**
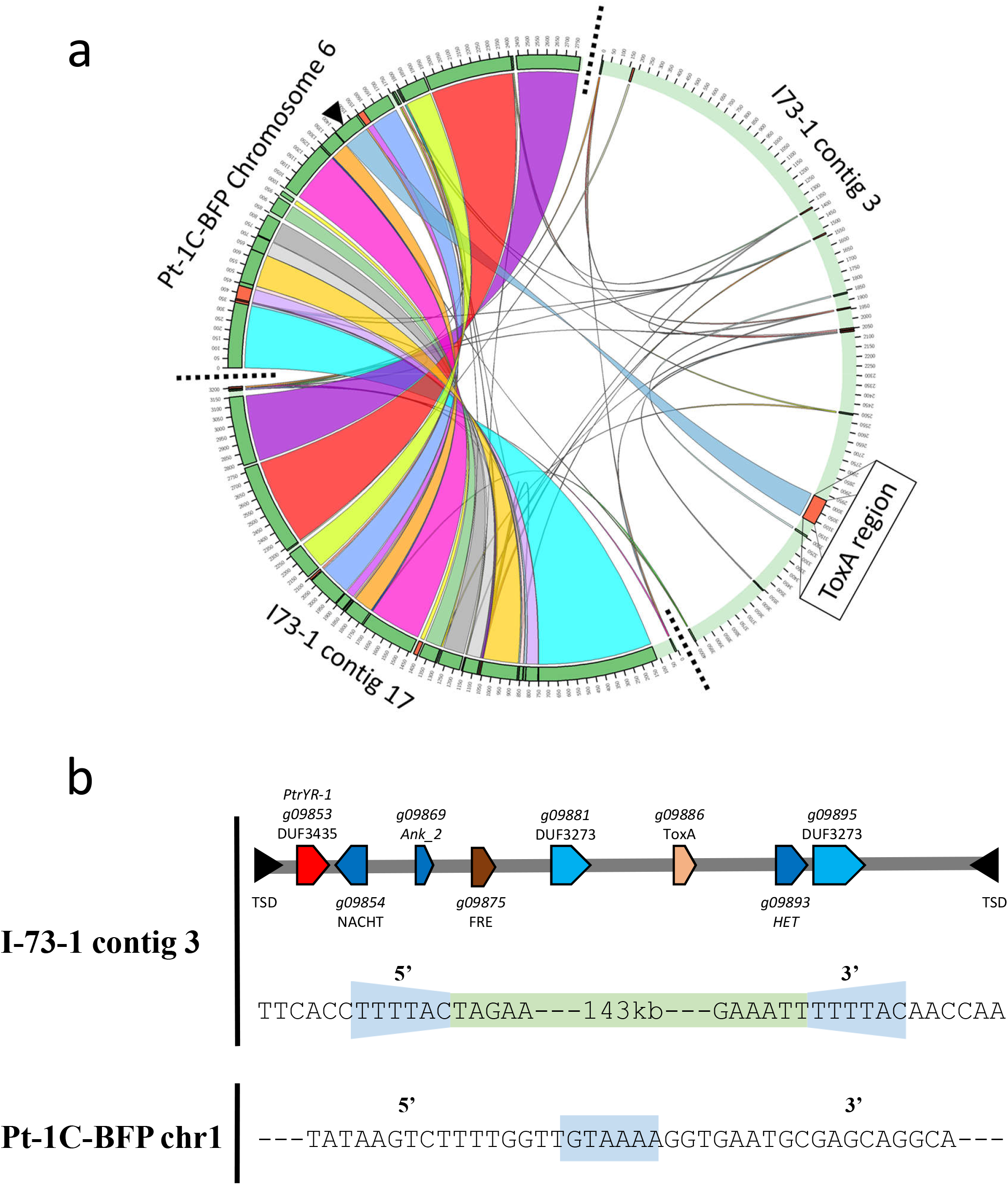
Evidence of translocation of *ToxA* containing region between two races of *Pyrenophora tritici-repentis*. **a** The *ToxA* containing chromosome 6 of isolate Pt-1C-BFP (race 1) is compared with the *ToxA* containing contig 3 and contig 17 of isolate I73-1 (race 8). Pt-1C-BFP chromosome 6 aligns almost completely to I73-1 contig 17, with the exception of the *ToxA* containing region, which aligns to contig 3. The majority of contig 3 aligns to Pt-1C-BFP chromosome 1 or chromosome 9 (Figure 3). **b** schematic of the ‘Horizon’ Starship class transposon.

Large-scale chromosomal rearrangements such as chromosome fusions and inversions were observed between I-73-1 and BFP. This was illustrated clearly by the chr 1 fragmentation in I-73-1, whereas chr 1 is the largest chromosome in BFP (10.2 Mb). A significant section of chr 1 aligned with contig 18 in I-73-1, but the remaining sections were found to be distributed between four contigs: 1, 3, 16, and 20. Additionally, BFP chr 8 was split between I-73-1 contigs 18 and 20. Approximately 30% of chr 3 (contig 2 in I-73-1) was inverted. Similar rearrangements on a smaller scale were observable in the alignment dotplot (Figure 3).

Similar to I-73-1, a chr 1 fragmentation was observed in D308. However, the fragmentation in the latter was less severe, with sections aligning to three contigs (D308 contigs 1, 2, and 15). There was a collinearity between D308 contigs 11 and 14 and chr 7 in BFP, and hence, these two contigs likely represent a single chromosome that would align fully with BFP chromosome 7 and I-73-1 contig 9. Furthermore, D308 contigs 7 and 25 fully aligned with BFP chr 2. The genome architectures of D308 and I-73-1 appear to exhibit higher collinearity between each other than to isolate BFP (Figure 3). We found similar types of rearrangements in alignments with M4 and DW-5 although they were fewer in number (not shown). There was no evidence for the presence of supernumerary chromosomes in I-73-1 or D308.

### ToxA is present on a Starship class transposon and ToxB is present within a massive putative transposon

*ToxA* in BFP was located on chr 6 (2.8 Mb), but in I-73-1, *ToxA* was carried on contig 3 (4.0 Mb) which aligns with BFP chr 1 and 9. While BFP chr 6 was present in I-73-1 as contig 17, the contig lacked *ToxA* and a 143 Kb segment (Figure 4A). This validates the previously reported translocation of *ToxA* in I-73-1 (13). Edge analysis and review of gene content within the 143 kb segment indicated target site duplications (or short direct repeats) at the edges and a tyrosine recombinase (gene_09853; DUF3435 domain) at the 5’ end, initially suggesting this element is a *crypton* (30). Recently, however, a new class of large mobile elements called *Starships* have been defined (31). An examination of other genes present within the transposon suggests that this element is actually a *Starship* with many similarities, including: the DUF3435 tyrosine recombinase ‘captain’, DUF3273 domains, ferric reductase, ankyrin repeats, heterokaryon incompatibility, NACHT nucleoside triphosphatase, and target site duplications (Figure 4B) (31). A full list of genes present are available in Supplemental File 2. The target site itself was identifiable in BFP chr1 (Figure 4B and Supplemental Figure 3), M4 (not shown), and D308 (not shown). We have named this new *Starship* ‘Horizon’. Additionally, an alignment of the ToxhAT transposon, which facilitated the movement of *ToxA* between Ptr, *Parastagonospora nodorum*, and *Bipolaris sorokiniana* (14), showed that this smaller 14 Kb TE is nested within Horizon.

Four copies of *ToxB* were found in the I-73-1 long-read assembly; each copy contained a single 49 bp exon. Three copies were located within a 12 Kb region on contig 12 (homologous to chr 4 in BFP), and the fourth copy was present on a small 6 Kb contig (contig 13), which failed to assemble with the chromosome-sized contigs. The Australian isolate M4 (race 1) offered a better scaffolding quality than BFP to examine the 12 Kb *ToxB* region, and the alignment of I 73-1 contig 12 to M4 contig NQIK01000005 indicated an even larger 294 Kb gap around the region of *ToxB* (Figure 5). Examination of the edges of this gap revealed the presence of terminal inverted repeats in I-73-1 and a potential target site duplication in M4. This same 294 Kb transposon is present in D308 on contig 9, and although the *ToxB* gene is missing in D308, its inactive homolog *toxb* is present within the same transposon as a single copy (Supplemental Figure 4A).

**Figure 5.**
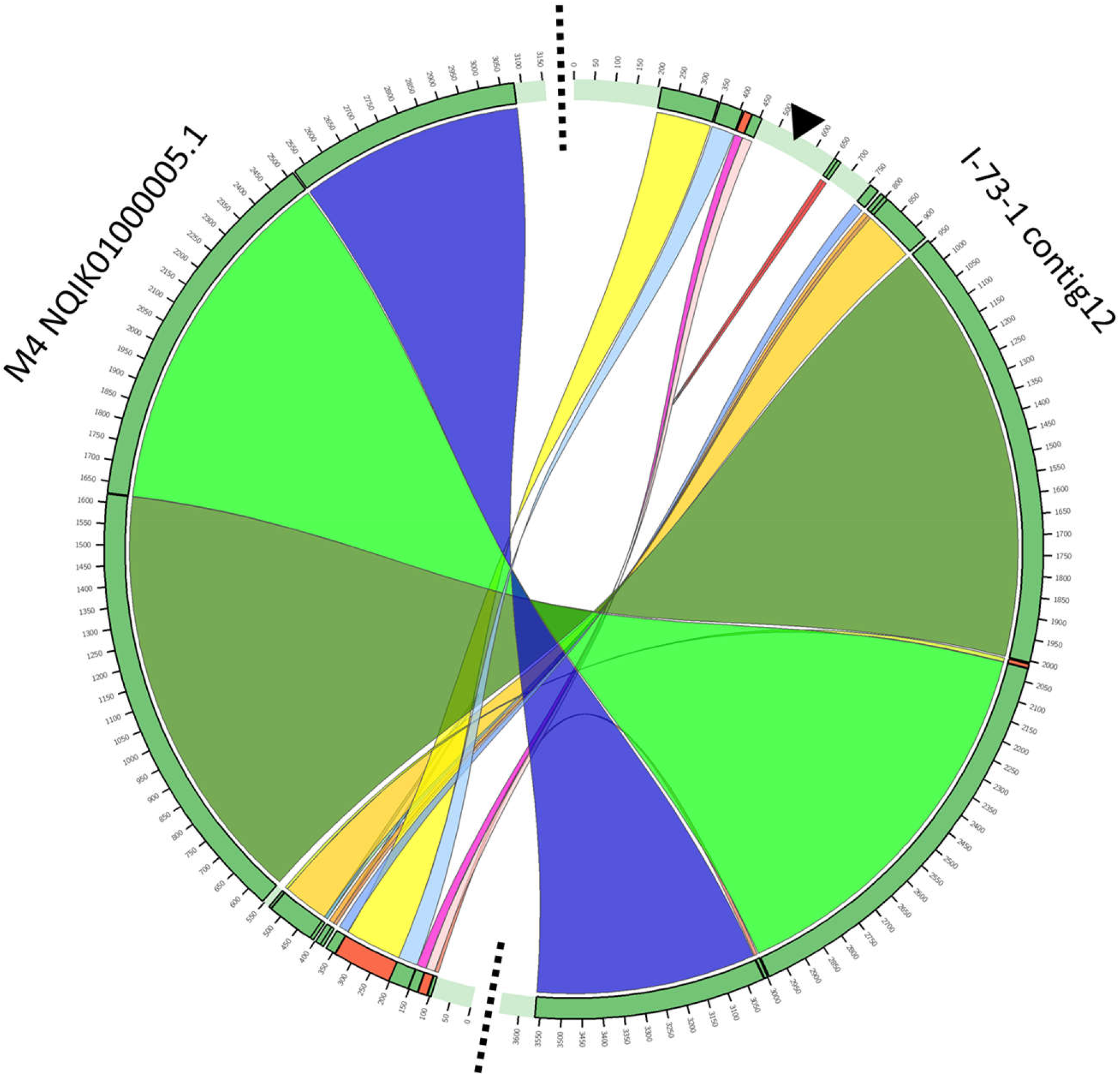
Circular alignment of contig 12 from race 8 isolates I-73-1 and contig NQIK01000005 from race 1 isolate M4. A large 294 Kb region which contains three copies of the *ToxB* (black arrow) is visible, this section did not align with any other contig of M4. The edges of the 294 Kb region revealed terminal inverted repeats suggesting transposon activity.

*ToxB/toxb* genes were found here to be located on an essential chromosome/contig in Ptr, as the same chromosome/contig is present in other isolates that do not carry the *ToxB* gene (BFP, D308 and M4). DW5 was reported to carry 10 copies of *ToxB* (32). Our BLAST output indicated 10 copies, but eight of these were on contigs ≤ 126 Kb, which may indicate misassembly. Only two copies of *ToxB* were on contigs of comparable size to chromosomes (CM025819 at 3.4 Mb and CM025824 at 2.2 Mb respectively). These contigs aligned with BFP chr 5 and 11 and I-73-1 contigs 7 and 16, respectively (Supplemental Figure 4B). The locations of *ToxB* copies in DW-5 do not appear to have homology to the putative transposon in I-73-1 and D308, which may indicate additional transposon activity associated with *ToxB*.

The chromosomes and contigs that possess *ToxB*/*toxb* were fully (or nearly fully) present in all isolates examined, indicating they are essential in nature or contain essential genes for Ptr. Additionally, the presence of core and accessory genes and their ratios in isolates I-73-1 and D308 were examined, and for I-73-1 contig 12, the ratio of core to accessory genes was 3.4, while for D308 contig 9 the ratio was 3.8, supporting the essential nature of these chromosomes (Supplemental File 3).

### Ptr exhibits a ‘one-compartment’ genome

The genome of many plant pathogens exhibits a two-compartment arrangement, one with AT-rich regions where effectors and TEs are embedded, and a second with GC-rich content (20). In order to evaluate if the Ptr genome exhibited compartmentalization, the intergenic distances (the number of nucleotides between genes) were measured and found to average ∼4,600 bp in both I-73-1 and D308. Density plots of the 5’ and 3’ intergenic distances showed a single hot spot at a similar size range (Figure 6). A scattering of genes in the upper right portion of the plot indicates that some genes are in gene-sparse regions, with intergenic distances as high as 100,000 bp in certain cases. The single hot spot (rather than two) is indicative of a ‘one-compartment’ genome, and although there are a number of genes in gene-sparse regions, their frequency does not support a ‘two-compartment’ genome. Additionally, analysis with RIPper showed that <2% of the I-73-1 and D308 genomes were affected by RIP, with <40 LRARs present in each.

**Figure 6.** Intergenic distances of all genes in *Pyrenophora tritici-repentis* of: **a** isolate D308; **b** isolate I-73-1. The 3’ intergenic length (x-axis) is the distance (bp) from the 3’ end of current gene to the 5’ end of next, and the 5’ intergenic length (y-axis) is the distance from the 3’ end of the previous gene to the 5’ end of the current gene.

### Ptr genome and expansion by transposable elements

Transposable element (TE) content in the short read assemblies ranged from 6.4% to 10.2% of the assembled genomes (Figure 7A and Supplemental File 4). The highest TE content occurred in isolates 92-171R5 (10.2%) and G9-4 (9.8%). A large proportion of the TEs in 92-171R5 were classified as ‘unknown’ repetitive elements. In G9-4, however, the increased number of TEs were primarily Class I-LTRs. The high TE content in these two isolates also correlates with larger genome sizes relative to the other assemblies (Figure 7B). TE content in 12 isolates (excluding 92-171R5 and G9-4) exceeded 7%, and the remaining 26 isolates have TE content ranging between 6.4% and 7.0%. All assemblies contained significant numbers of ‘unknown’ repetitive elements. The program EDTA (Extensive de-novo TE Annotator) uses structural features to identify intact TEs at the beginning, and then classifies them into families based on coding features. RepeatModeler2 was used to identify repeats not initially reported, but due to a lack of homology and coding features, they were labelled as ‘unknown’ TEs.

**Figure 7.** Transposable element content and genome size variation of *Pyrenophora tritici-repentis* from short-read assemblies. **a** Contribution of TEs (%) to total genome size across 40 isolates. **b** Contribution of each TE class to genome size for individual isolates.

The long-read assemblies of isolates I-73-1 and D308 showed a significantly larger number of transposable elements (>150%). TEs represented 6.8% and >18.3% for the short- and long-read assemblies, respectively. In both isolates, the increased TE content was due to a greater incidence of Class I transposons, primarily Copia and Gypsy elements. Additionally, many previously uncategorized transposons (labelled as ‘unknown’ in the short-reads), were reclassified with accurate transposon labels. Almost all groups of transposons identified in the short-read assemblies showed some increased incidence in the long-read assemblies, with the exceptions of the Tc1-Mariner, Helitrons, and PIF-Harbinger classes, which were consistent between assembly types.

## DISCUSSION

In this study, we performed a global pangenome analysis of Ptr, an important foliar pathogen of wheat. In total, 41 Ptr genomes carrying various combinations of all known and unknown effectors were analysed. These isolates represent various geographical origins, extending from regions where wheat is of relatively recent introduction, to regions encompassing the centre of wheat origin. We identified major rearrangements at the chromosomal level among five assembled Ptr genomes, two of which were generated in this study (I-73-1 and D308) and three of which were previously described (BFP, M4, and DW5). These rearrangements included chromosomal fusions, segment inversions, and translocations. We showed that the Ptr genome appears to tolerate large-scale structural reorganizations, genome size variation, and has remarkable plasticity. Previously, a worldwide collection of Ptr isolates, some of which were included in this study, had shown independent chromosomal location of the virulence genes *ToxA* and *ToxB* and extensive genome plasticity (karyotypes), where aneuploidy, proliferation of repetitive DNA and transposon activities were suggested as driving mechanisms (13). Here, we showed plasticity in the Ptr genome is likely to be facilitated by the proliferation of TEs and the expansion of the accessory genome, in addition to the role of transposable elements in virulence gene shuffling.

Fungi are known to display high variability in their genomes, and the ability of plant pathogenic species to gain virulence genes embedded in a “pathogenicity island” and carried on supernumerary chromosomes that transfer horizontally as whole or in part is well described (33). In recent years, more reports have emerged on the ability of virulence genes to transfer on large transposons among plant pathogens (31). Overall, the ability of pathogenic fungi to evolve rapidly to invade their hosts or cause outbreaks is illustrated through their ability to recombine, mutate and shuffle their genetic components either vertically or horizontally. Traditionally, a ‘two-compartment’ genome has been associated with a ‘two-speed’ genome in pathogenic fungi. In this case, Ptr appears to possess a ‘one-compartment’ genome; however, this does not preclude the species from having a ‘two-speed’ genome driven by copy number variation and TEs not associated with compartmentalization (20), as we know *ToxB* is present as multiple copies in Ptr. Other effectors or genes associated with virulence may be present in multiple copies, but this remains to be explored.

### The intraspecies translocation of ToxA in Ptr is facilitated by the Starship transposon ‘Horizon’

*ToxA* is present in other fungal species such as *Parastagonospora nodorum* and *Bipolaris sorokiniana* (7, 14). The independent horizontal transfers between these species has been explored in detail, with the transfer being facilitated via a hAT transposon dubbed ToxhAT, which is 14 Kb in size (14). Previously, *ToxA* was found to be located on the same essential chromosome in all tested Ptr isolates with the exception of I-73-1, where it was shown to be translocated to a larger non-homologous chromosome, as indicated by pulse-field gel electrophoresis followed by southern hybridization (13). Although ToxhAT is present, we validate the translocation of *ToxA* in I-73-1 via ToxhAT, but nested on a much larger putative 143 Kb *Starship* transposon, which we have named ‘Horizon’. This mobile element was integrated into an essential chromosome (contig 3; 4Mb) in I-73-1, corresponding to chr 9 in BFP. The *ToxA* coding gene in all Ptr isolates tested here has a very conserved sequence belonging to one haplotype (PtrH1) (34) (data not shown), and this supports the hypothesis of its recent integration into the Ptr species (7, 35).

The movement of *ToxA* in different types of mobile elements in Ptr and the nesting of transposons is indicative of this pathogen’s ability to evolve and acquire virulence rapidly. The nesting of mobile elements with virulence factors has been linked to the evolution of pathogenicity in other species as well (14, 36, 37) and the association of TEs with effectors has been well documented in other fungal pathogens, most recently in another wheat pathogen *Zymospetoria tritici*, which is consistent with our findings (38). Intrachromosomal translocation of *ToxA* was also described in *B. sorokinaina* (14), and highlights the mobility of *ToxA* within Ptr and *B. sorokinaina* via transposon activity and/or genomic recombination. Multiple HGT events were previously suggested for ToxA and its 14 Kb surrounding region, the ToxhAT transposon (14). In this study we showed the nesting of ToxhAT in a larger mobile element Horizon which was not present in other ToxA containing isolates like BFP and M4. These findings add support to the idea that ToxhAT may have been transferred to Ptr multiple times, with one HGT event nesting ToxhAT within Horizon. Additionally, were Horizon to share sequence homology with other isolates or species in the future that would also support the hypothesis of multiple *ToxA* HGT events.

### The ToxB gene is embedded on a large putative transposon in Ptr

For the first time, we revealed the possible movement of *ToxB* as a multi-copy gene clustered in race 8 isolate I-73-1 (contig 12), and the single copy *toxb* homolog on contig 5 in D308 race 3 on a large 294 Kb transposon potentially similar to that of *ToxA*, although an analysis that confirms this classification is required. No clear evidence of *ToxB* horizontal transfer has been found to date, but homologs of *ToxB* are present in closely related species (e.g., *P. bromi, Cochilobolus sativus, Alternaria alternata, Magnaporthe grisea*) (8). It has been suggested that *ToxB* was acquired vertically from a common ascomycete ancestor (8, 15). However, our discovery of *ToxB/toxb* on a large putative transposon inserted into an essential chromosome provides the new possibility of an alternative mechanism of acquisition via a horizontal transfer event older than that of *ToxA* (given the higher variability in *ToxB*/*toxb* reported sequences). Moreover, our analysis suggested a potential transposon(s) associated with *ToxB* in the previously published isolate DW-5, which provides further evidence of potential horizontal mobility of *ToxB*.

### CAZymes are essential part of the Ptr genome

Ptr is a necrotrophic pathogen that posses the ability to directly penetrate the host epidermal cells soon after spore germination (39), and this penetration is likely facilitated by the fungus ability to secrete cell wall degrading enzymes. The CAZymes and CAZy families identified in the Ptr pangenome support that Ptr is adapted for plant cell wall degradation, as >30% of the total CAZome belongs to families involved in dismantling structural polysaccharides, such as cellulose, hemicelluloses, and the plant cuticle. Phytopathogenic fungi are known to deploy a number of CAZymes for invasion and infection (26, 28, 29). As an example, *Fusarium culmorum* has been shown to degrade cellulose, xylans, and pectins during invasion of wheat spike tissues (40). While previous work has linked the relatedness of CAZymes to fungal taxonomy (41), it is not known if there is functional specialization in different Ptr isolates. CAZomes may be tailored to accommodate unique structures in cell wall polysaccharides of different wheat varieties or plants growing in different geographic regions. Indeed, the diversification of AA9 structure, sequence, and expression levels in isolates of the phytopathogen *Rhizoctonia solani* has been proposed to be essential for the differential pathogenicity of these strains in rice and soybean (42), and the same may be true for Ptr and its hosts.

### Ptr has an open pangenome with a high accessory gene content

In an open pangenome, each added genome increases the number of accessory genes, while the number of shared (core) genes decreases (43). In this work, we have shown that Ptr has an open pangenome. We also showed that Ptr, unlike other ascomycetes, contains a large proportion of accessory genes (57%) with a relatively small core gene set (43%). This is a significantly smaller core than the estimated 69% core previously reported in the pangenome of 11 Ptr isolates (17), and the 60% core estimated in the pangenome of the wheat pathogen *Z. tritici*, which was based on 19 isolates (44).

It has been suggested that plant pathogens possess accessory genomic elements related to pathogenicity (45). However, we showed that the Ptr pangenome possesses a higher number of accessory genes in the non-pathogenic and weakly virulent isolates; these isolates also exhibited a larger proportion of orphan (singleton) genes. Approximately 70% of singleton genes had unknown functions, but perhaps these genes are involved in adaptation for a lifestyle divergent to wheat pathogenesis (23). The open pangenome of Ptr may explain its ability to adapt to a wide variety of hosts, geographical regions, and environmental conditions. Despite its homothallic mating style, this pathogen evidently acquired virulence by horizontal transfer of a large segment of DNA from other fungal species (7, 14). The core genome is usually protected from high recombination and mutation rates in order to preserve its essential biological function, and the accessory genome is assumed to be under rapid evolution (45). It is still unclear how the accessory genome in Ptr evolves and why the non-pathogenic isolates exhibited higher contents of accessory genes and TEs.

The observed separation of isolates based on ToxA production in the core protein phylogeny and the accessory protein hierarchical sets may be due to the host specialization of Ptr between bread and durum wheat. A similar observation has been made before, using simple sequence repeats on a similar collection of isolates (12). The non-pathogenic isolates (except T-126-1 from Tunisia) were also distantly related to the rest of the isolates, and we observed a divergence in TE and effector content in these isolates, which is also an indicator of host specialization.

We also found variation in the genome size, particularly between pathogenic and non-pathogenic isolates. The genome size in the Dothideomycetes, based on the analysis of 101 species, varies 10-fold: from less than 17 Mb to more than 177 MB (46). Our Ptr genome size based on the more accurate long read assemblies averaged at 39.8 Mb, which was very consistent with previous genome size estimates for long-read Ptr in previous studies (17, 19, 32). However, we showed that the non-pathogenic race 4 and weakly virulent race 5 isolates, G9-4 and 92-171R5 genome size were ∼2 Mb larger than the average pathogenic isolates, and both isolates have an average of 3.1% more TE content than the other isolates. This clearly indicates an expansion of TEs in the genomes of these isolates. This larger genome size in non-pathogenic isolates was not expected, as previous reports of Ptr genome sizes indicated that pathogenic isolates have larger genomes, which was not consistent with our observations (17). The larger genomes reported in pathogenic races/species relative to their non-pathogenic counterparts has been attributed to the presence of more repetitive elements, and it was posited that the need of pathogenic species to evolve in an arm’s race with its host could explain such genome expansion (47). Indeed, in some plant pathogens the acquisition of an entire supernumerary chromosome, often rich with repetitive sequences, by horizontal transfer is evident (48). However, not all pathogenic species follow this trend of a larger genome (35). A reduction in pathogen genome size may be an evolutionary mechanism to aid adaptation to specific environments or other niches, and could explain the recent specialization of pathogenic species from related generalist non-pathogens (49–52). Many pathogens adapt to exploit a limited number of species efficiently, and in some species, even a limited number of genotypes within a species (39) with rapid gene gains and losses previously linked to virulence adaptation in other studies (53, 54).

Nonetheless, while a larger genome size was found in the non-pathogenic race 4 isolate G9-4 and the weakly virulent race 5 isolate 92-171R5, this was not the case for the other race 4 isolates, 90-2 and T126-1, which had genome sizes similar to the pathogenic strains. As such, it is not easy to make a clear conclusion regarding genome size and pathogenicity in Ptr, and there is a need to explore additional non-pathogenic genomes. In previous studies, comparisons with Ptr non-pathogenic genomes were based on a single race 4 isolate (SD20-NP) (16, 17). Additionally, in many other ascomycetes, the number of non-pathogenic genomes studied is limited in comparison with pathogenic genomes (55). It is worth noting that non-pathogenicity in Ptr often has been assumed based an isolate’s inability to cause disease symptoms on a limited number of host genotypes. However, what were assumed to be non-pathogenic race 4 isolates were recently found to cause extensive necrosis on specific durum wheat genotypes (56).

### Conclusions

Collectively, this work highlights the high plasticity and potential adaptability of Ptr as a global wheat pathogen. The large-scale chromosomal rearrangements, open pangenome, and the extensive accessory gene and TEs content of its genome reflect its cosmopolitan nature. In addition, the nesting of the virulence genes *ToxA* and *ToxB* within multiple transposon types is a significant indication of the rapid evolutionary nature of this pathogen.

## Methods and Materials

### Isolates, DNA extraction, and sequencing

Isolate details are provided in Table 1. Virulence phenotypes were previously confirmed (4, 12, 13, 57). Isolates collected before 2016 and received from other labs were subjected to single spore isolates and their phenotypes confirmed on a wheat differential set as described by Aboukhaddour et al., (2013). Isolates were grown in 100 mL 1/4 concentration PDB and cultures incubated in a shaker (100 RPM) at room temperature (∼25°C), without light, until the fungal mat covered the surface of the medium. Fungal mats were harvested and washed twice using autoclaved mili Q water. Mats were placed in whirl pack bags on dry ice until freeze-dried. Dried mats were stored at −20°C until DNA extraction (no longer than 14 days). Genomic DNA was extracted using a ‘Genomic-tip 20/G’ kit (Qiagen) and sequenced with 150 bp paired-end, 400 bp inserts at 100x coverage with Illumina HiSeq X. DNA from I-73-1 and D308 was extracted using a ‘Genomic-tip 100/G’ kit (Qiagen) and long-reads sequenced with PacBio RS II at 100x coverage. All sequencing was performed by the Centre d’expertise et de services Génome Québec (Montreal, Canada). The reference isolate BFP (accession GCA_000149985) (16) was included in the pangenome analysis and used for full genome alignments. Isolates M4 (GCA_003171515) (17) and DW5 (GCA_003231415) (32) were also included for *Tox* gene and transposon analysis.

### De novo assemblies

Read quality was assessed with FASTQC (58) and poor quality reads filtered. Reads were filtered using the standard Kraken2 database (59) and any reads tagged as non-fungi were omitted. A subset of isolates (90-2, AB88-2, ASC1, AZ35-5, and I-72-1) were selected to test assembly program suitability using the Illumina reads. The assemblers were Shovill with SPAdes (60, 61), Shovill with MEGAHIT (61, 62), SOAPDenovo2 (63), and CLC Genomics Workbench 12 (Qiagen) all program arguments available in GitHub repository. QUAST (64) output and BUSCO scores were used to assess assembly quality. All remaining short-reads were assembled with Shovill/SPAdes. Long-reads were assembled with Flye (65) and polished with short-read data using Pilon (66). Completeness of the long-read assemblies was also assessed by the alignment of raw reads (Supplemental Figure 5).

### Gene annotations

FunGAP (v1.0.1) is an annotation pipeline specifically for fungi (67). FunGAP makes use of many programs: RepeatModeler (68), RepeatMasker (69), HISAT (70), Trinity (71), Augustus (72), MAKER (73), BRAKER (74), InterProScan (75), and BLAST+ (76). MAKER and BRAKER utilize RNA-seq reads. RNA-seq reads from vegetative Ptr mycelia were retrieved from BioPlatforms Data Portal (https://data.bioplatforms.com/; Sample ID: 102.100.100.14350) (17). FunGAP provides a script to select a prediction model for Augustus, with *Botrytis cinerea* selected as the model in this case. Default settings provided by FunGAP were used to annotate all assemblies. After pangenome analysis (discussed below), representatives from the core and accessory gene clusters were used for functional annotation with Pfam v28.0 (77). All predicted gene sets were assessed for completeness using the Ascomycota BUSCO gene set (odb9) (78, 79).

### Pangenome analysis

Pangloss is a pangenome pipeline designed for microbial eukaryotes like fungi (21). Panoct (80) is the primary program for calculating a pangenome. A custom bash script was used to modify *.gff3 files from FunGAP into *.attributes format required by PanOCT. Output from all-vs-all BLASTp (E10^−4^) was used to determine if a gene belongs in the core or accessory genome. Pangloss does not distinguish singletons from the accessory genome, a custom bash script separated singletons. Binary gene presence or absence tables were parsed to generate figures showing the percentage of the genome comprised by core, accessory or singleton genes, total genes in the pangenome as new genomes were added, and the number of genes in accessory clusters. Statistics including t-tests, ANOVA and Tukey’s HSD were performed with R (v3.4.3) in RStudio.

### CAZymes and effectors

Phobius v1.01 (with --short) (81) was used to filter amino acid sequences for signal peptides and transmembrane domains. Sequences with signal peptides and without transmembrane domains were used as input for EffectorP-2.0 (82). Genes identified as potential effectors (probability >50%) were extracted from gene sets and used as input for Pangloss to assess presence/absence between isolates. Predicted protein sequences produced by FunGap from all isolates were annotated by dbCAN2 (83) and manually curated for selected GH, CE, PL, and AA families relevant for plant cell wall degradation from the CAZy database (24).

### Phylogeny and accessory sets

Individual alignments for each of the 10,159 amino acid core orthologue clusters was performed using MUSCLE (v3.6) (84). Bash and python scripts were used to concatenate and combine the aligned core amino acid gene set for each isolate creating a single alignment. The core alignment was input for RAxML (85) to generate a Maximum Likelihood (ML) phylogeny using the PROTGAMMA model, 1,000 bootstrap replicates, and a starting seed of 10. Variant call files were generated via GATK HaplotypeCaller (86) using BFP as a reference. SNPs were converted to fasta format with vcf2phylip (87) and input into RAxML for a ML tree using GRTCAT model. The ML trees were visualized with FigTree (88) and the Tox/location content added manually. Binary data was filtered to include only accessory genes and was input for the R package HierarchicalSets with default settings (89).

### Transposable element content

Transposable elements (TEs) were identified and categorized using EDTA with the higher sensitivity setting (--sensitive 1) (90), which utilizes several programs: LTRharvest (91), LTR_FINDER (92), LTR_FINDER_parrallel (93), LTR_retriever (94), and TIR-Learner (95), Generic Repeat Finder (96), HelitronScanner (97), and TEsorter (98). Output was aggregated and the total TE content per genome (stacked histogram) and TE content as a function of genome size (Mb) visualized.

### Genome organization

Pair-wise full genome alignments of long-read assemblies: I-73-1, D308, BFP, M4, and DW-5 were performed with minimap2 (99) and visualized with dotPlotly (100). For chromosomes with *ToxA* and *ToxB* putative transposons, syntenic blocks with a minimum length of 6,500 bp were generated using Sibelia (v3.0.7)(101), then visualized using Circos (v0.69-8)(102). Selected proteins within the putative transposons were modelled with Phyre2 (103). Intergenic distances were calculated from gff3 files using an adapted R-script (104) and plotted using an hexagonal heatmap of 2d bin counts geom_hex with 50 bins. Long-read genomes were uploaded to RIPper (105) to assess RIP and LRAR content. The core vs. accessory nature of contigs was assessed by alignment (i.e., presence in all isolates) and evaluation of the ratio of core to accessory genes, percentage of total genes, and genes per Mb on each contig.

### Data availability

Isolate accession numbers are available in file Supplemental File 5. Raw data is available under NCBI BioProject PRJNA80319. Scripts and program settings used in this work are available at https://github.com/fungal-spore/Ptr-pangenome-paper.

## AUTHOR CONTRIBUTIONS

RG performed most of the analyses including assemblies/annotations, pangenome, phylogenetics, TE content, chromosomal organization, effectors, and *ToxA* and *ToxB* transposon analysis. The first draft of this manuscript was prepared by RG, RA, and MM. MM aided and guided the transposon analysis, and project guidance. ROP aided with assemblies and analysis guidance. MH performed DNA extractions. KEL and DWA performed CAZyme work. SES and FD provided isolates and input. RA conceived/guided project, performed single-spore isolation for a number of isolates and confirmed phenotyping. All authors contributed to editing and improving the manuscript.

## DEDICATION

This work is dedicated to the memory of the late Dr. Lakhdar Lamari, a passionate scientist and an exceptional teacher who had made significant contributions to the tan spot research community and the world of necrotrophic pathogens. Many of the isolates included in this study were collected by him during his trips to the Fertile Crescent, North Africa, and Caucuses regions.

## ACKNOWLEDGEMENTS

We would like to thank AAFC BioCluster Team, Therese Despins (culture maintenance), and Kathryn Wilde (image formatting), Sana Kamel and Mejda Cherif (University of Carthage, Tunis; Tunisian isolates). Funding: Agriculture and Agri-Food Canada and Alberta Wheat Commission and Saskatchewan Wheat Development to RA. The funding bodies were not involved in the design of the experiments and collection, analysis, and interpretation of data, or in the writing of this manuscript.

## Supplemental Figure Captions

**S01**. Genes were clustered based on the number of isolates in which they were present (e.g. genes in cluster 2 are present in two isolates, genes in cluster 3 are present in three isolates, etc.). Genes in low clusters may represent recently gained genes, as only a few isolate contain them, while genes in high clusters may represent recently lost genes, as most isolates contain them. Clusters 1 and 41 were omitted as they represent singletons and the core gene set respectively.

**S02**. Maximum likelihood phylogenies created by RAxML based on SNP data. **a** ML tree with all isolates; **b** ML tree with the divergent outgroup omitted to aid reading of the other branches.

**S03**. Truncated MAUVE alignment of BFP chr1 and I-73-1 contig 3 which shows the target insertion site and short direct repeats of putative *ToxA* transposon.

**S04**. Circular alignment of *ToxB* carrying contigs. **a** contig 5 from race 3 isolate D308, contig 12 from race 8 isolates I-73-1, and contig NQIK01000005 from race 1 isolates M4. A large 294 Kb region which contains three copies of the *ToxB* (black arrow) is visible which aligns with a section in D308 which contains a single copy of the inactive *toxb* (grey arrow). **b** DW-5 contigs (CM025819.1 and CM025824.1) to matching Pt-1C-BFP chromosomes (chr 5 and 11 respectively). Sections containing *ToxB* (black arrows) do not appear to share homology with each other or the reference chromosomes indicating possible transposon activity.

**S05**. Alignment of raw read data to long-read assemblies; **a** I-73-1; **b** D308.

## Supplemental Files

**SF01**. Curated output from dbCAN for grouping and counting the number of genes in specific CAZyme families.

**SF02**. Annotation of genes present within the 143 kb *Starship* transposon ‘Horizon’ present in the race 8 isolate I-73-1.

**SF03**. Number of core, accessory, and singleton genes on individual contigs of the long-read assemblies of I-73-1 and D308.

**SF04**. Table summary of transposable element content for each isolate of Ptr.

**SF05**. List of GenBank accession numbers associated with the genomes generated by this study and of the genomes downloaded and used for comparison.

