## Supplemental Figure 1 for "Dissecting the *Pyrenophora tritici-repentis* (tan spot of wheat) pangenome"

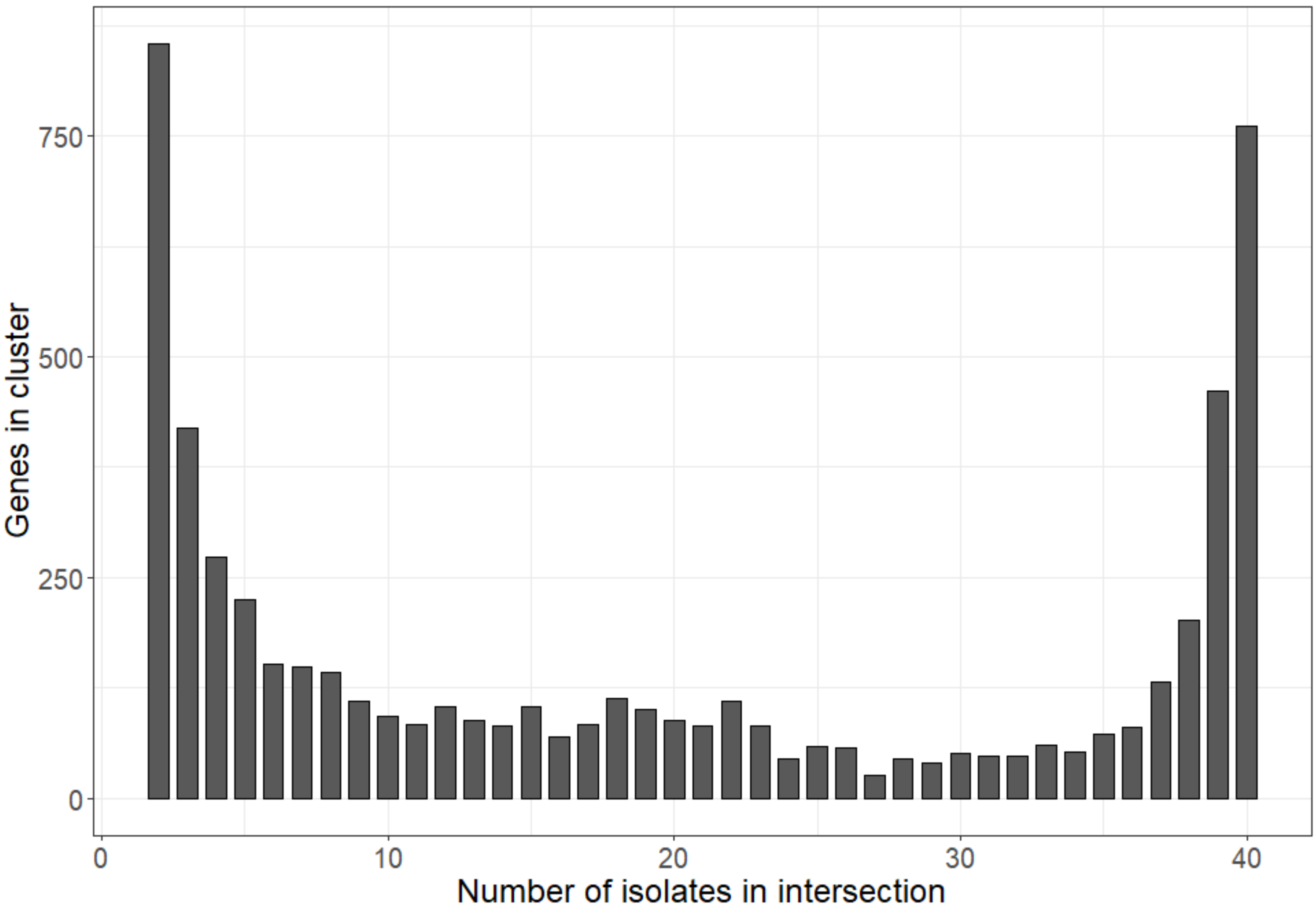


**S01.** Genes in Ptr were clustered based on the number of isolates in which they were present (e.g. genes in cluster 2 are present in two isolates, genes in cluster 3 are present in three isolates, etc.) Genes in low clusters represent recently gained genes, as only a few isolates contain them, while genes in hig clusters represent recently lost genes, as most isolates contain them. Cluster 1 and 41 were omitted as they represnt singletons and core genes respectively.
