## Supplemental Figure 2 for "Dissecting the *Pyrenophora tritici-repentis* (tan spot of wheat) pangenome"

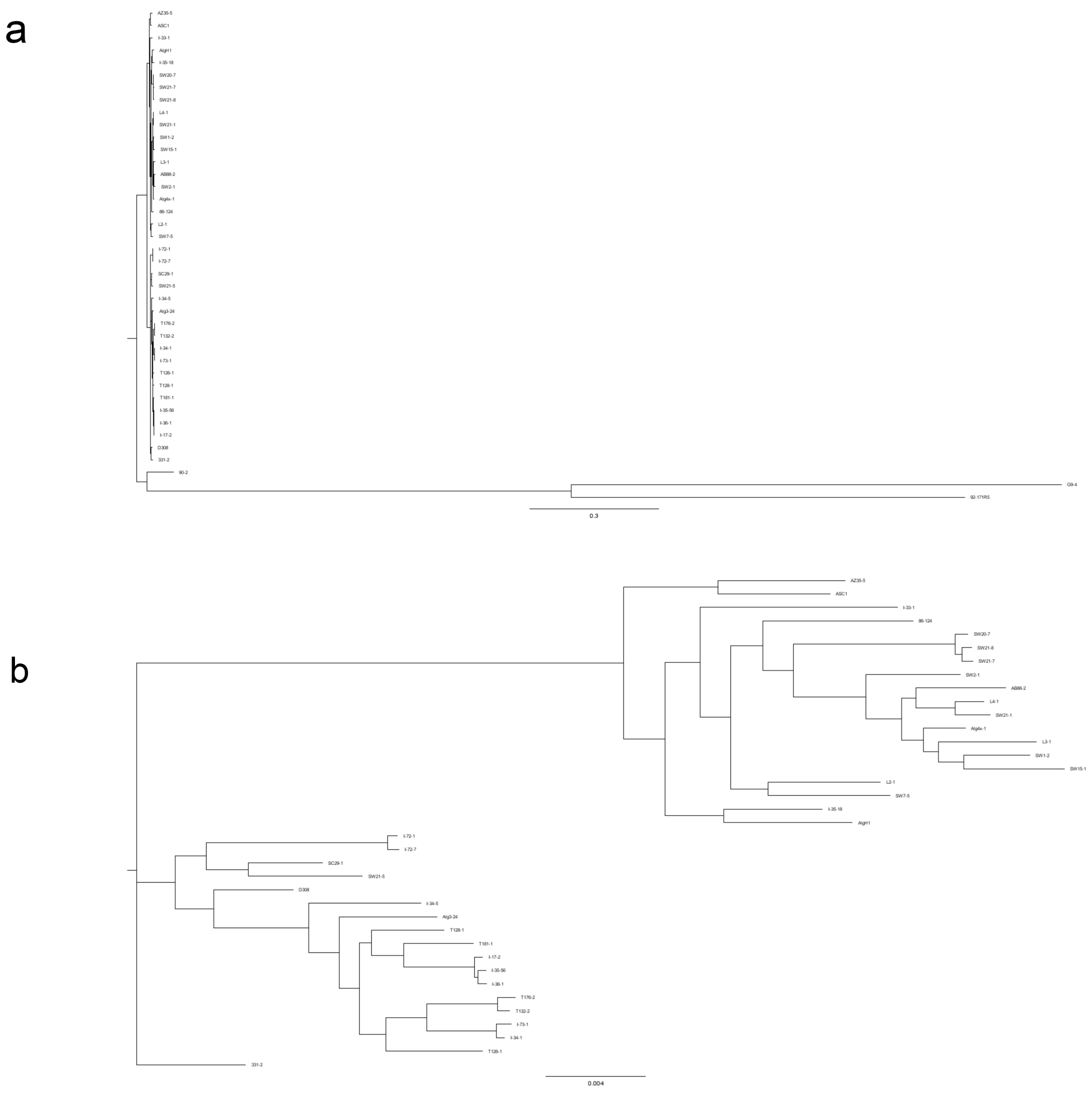


**S02.** Maximum likelihood phylogenies created by RAxML based on SNP. **a** ML tree with all isolates, branching is nearly identical to the core protein ML tree in Figure 2; **b** ML tree with the divergent outgroup (90-2, G9-4, and 92-171R5) omitted to aid reading of the other branches.
