## Supplemental Figure 3 for "Dissecting the *Pyrenophora tritici-repentis* (tan spot of wheat) pangenome"

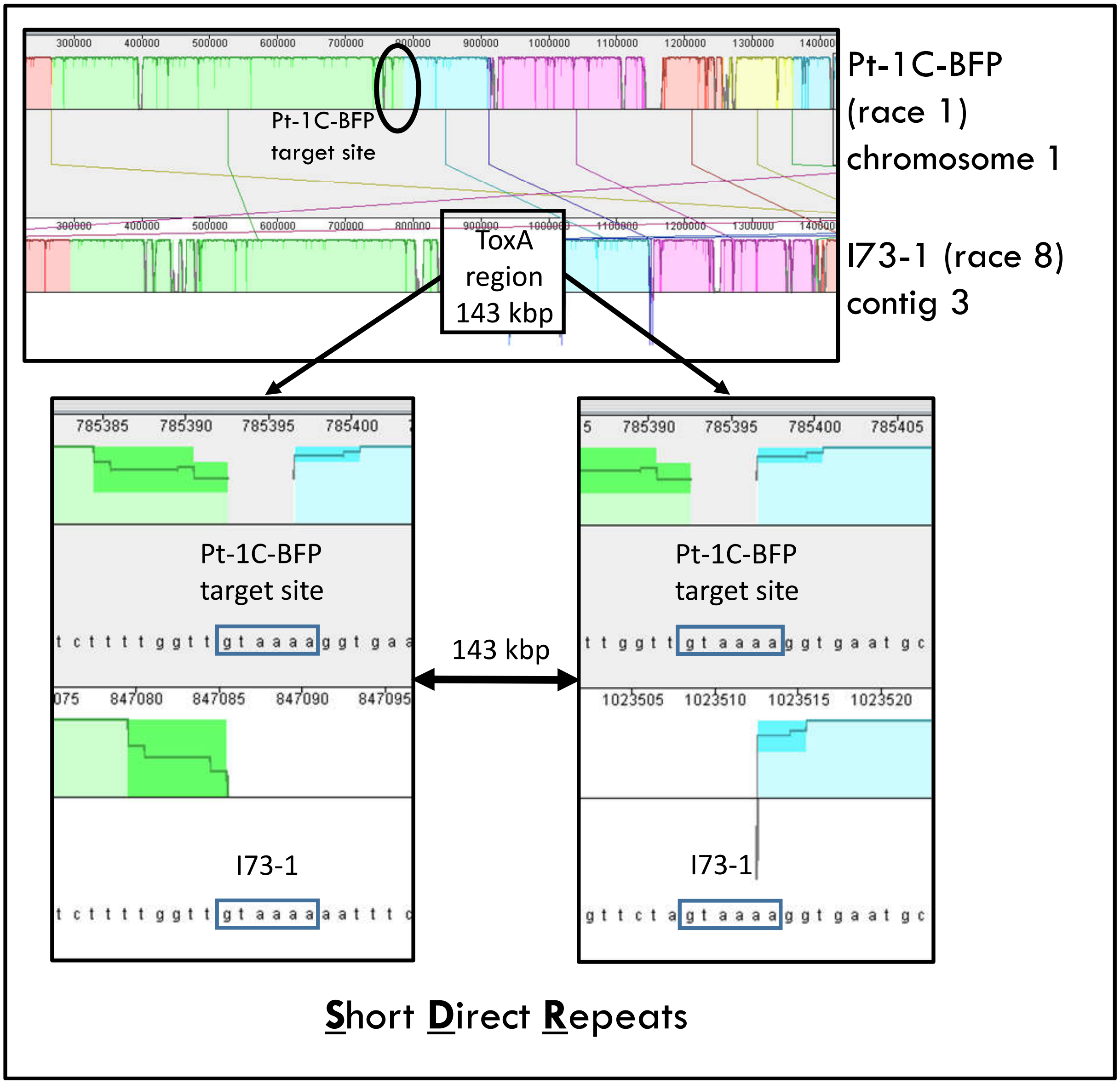


**S06.** Truncated MAUVE alignment of BFP chr1 and I-73-1 contig 3 which shows the target insertion site and short direct repeats of the *ToxA* transposon Horizon.
