## Supplemental Figure 4 for "Dissecting the *Pyrenophora tritici-repentis* (tan spot of wheat) pangenome"

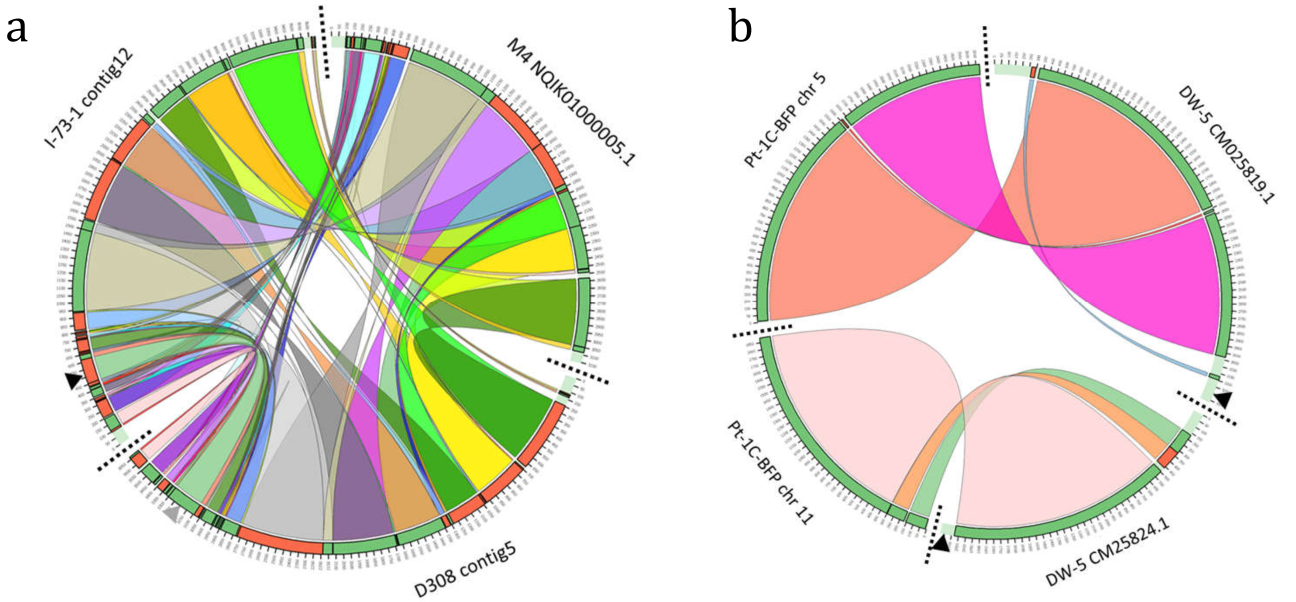


**S04.** Circular alignment of *ToxB* carrying contigs. **a** contig 5 from race 3 isolate D308, contig 12 from race 8 isolates I-73-1, and contig NQIK01000005 from race 1 isolates M4. A large 294 Kb region which contains three copies of the *ToxB* (black arrow) is visible which aligns with a section in D308 which contains a single copy of the inactive *toxb* (grey arrow). **b** DW-5 contigs (CM025819.1 and CM025824.1) to matching Pt-1C-BFP chromosomes (chr 5 and 11 respectively). Sections containing *ToxB* (black arrows) do not appear to share homology with each other or the reference chromosomes indicating possible transposon activity.
