## Supplementary figures and images for "Dissecting the *Pyrenophora tritici-repentis* (tan spot of wheat) pangenome"

### Supplemental Figure 5

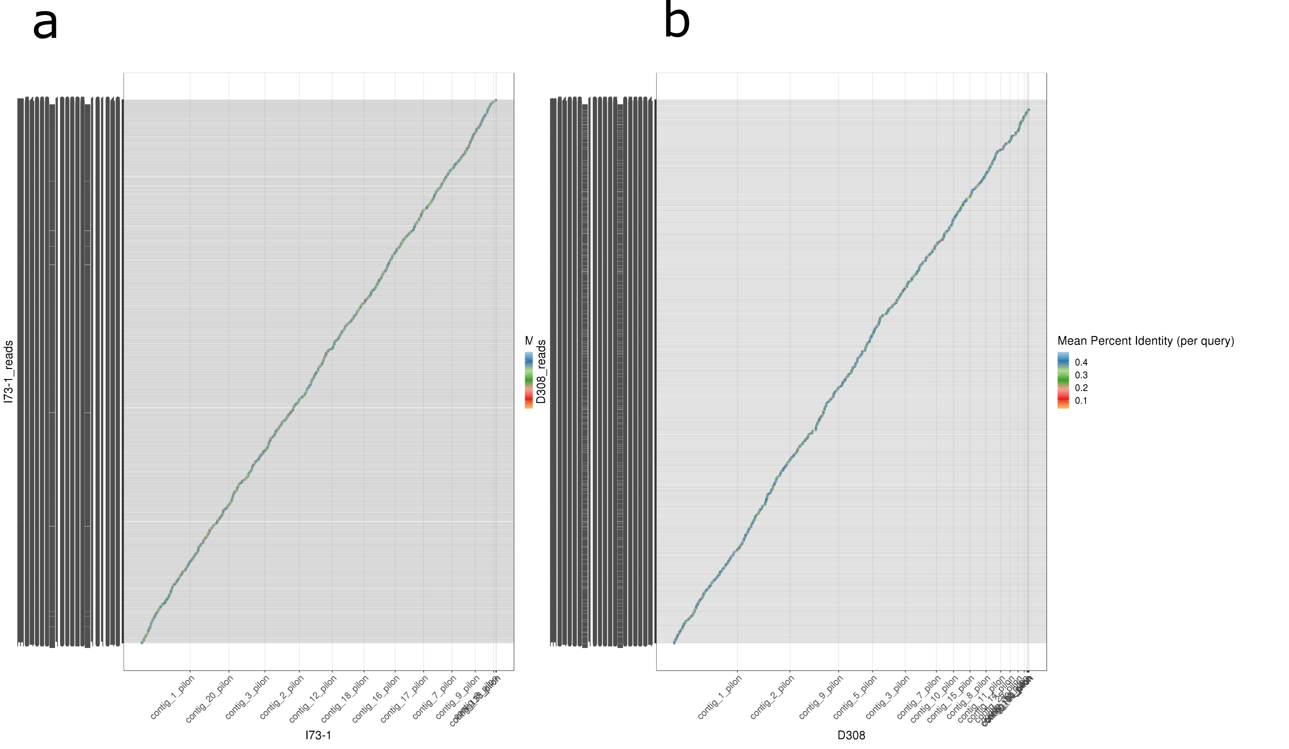


**S12.** Alignment of raw read data to the long-read assemblies of: **a** I-73-1; **b** D308
